# Innovations in Primate Interneuron Repertoire

**DOI:** 10.1101/709501

**Authors:** Fenna M. Krienen, Melissa Goldman, Qiangge Zhang, Ricardo del Rosario, Marta Florio, Robert Machold, Arpiar Saunders, Kirsten Levandowski, Heather Zaniewski, Benjamin Schuman, Carolyn Wu, Alyssa Lutservitz, Christopher D. Mullally, Nora Reed, Elizabeth Bien, Laura Bortolin, Marian Fernandez-Otero, Jessica Lin, Alec Wysoker, James Nemesh, David Kulp, Monika Burns, Victor Tkachev, Richard Smith, Christopher A. Walsh, Jordane Dimidschstein, Bernardo Rudy, Leslie Kean, Sabina Berretta, Gord Fishell, Guoping Feng, Steven A. McCarroll

## Abstract

Primates and rodents, which descended from a common ancestor more than 90 million years ago, exhibit profound differences in behavior and cognitive capacity. Modifications, specializations, and innovations to brain cell types may have occurred along each lineage. We used Drop-seq to profile RNA expression in more than 184,000 individual telencephalic interneurons from humans, macaques, marmosets, and mice. Conserved interneuron types varied significantly in abundance and RNA expression between mice and primates, but varied much more modestly among primates. In adult primates, the expression patterns of dozens of genes exhibited spatial expression gradients among neocortical interneurons, suggesting that adult neocortical interneurons are imprinted by their local cortical context. In addition, we found that an interneuron type previously associated with the mouse hippocampus—the “ivy cell”, which has neurogliaform characteristics—has become abundant across the neocortex of humans, macaques, and marmosets. The most striking innovation was subcortical: we identified an abundant striatal interneuron type in primates that had no molecularly homologous cell population in mouse striatum, cortex, thalamus, or hippocampus. These interneurons, which expressed a unique combination of transcription factors, receptors, and neuropeptides, including the neuropeptide *TAC3*, constituted almost 30% of striatal interneurons in marmosets and humans. Understanding how gene and cell-type attributes changed or persisted over the evolutionary divergence of primates and rodents will guide the choice of models for human brain disorders and mutations and help to identify the cellular substrates of expanded cognition in humans and other primates.

## INTRODUCTION

Vertebrate brains contain many specialized brain structures, each with its own evolutionary history. For example, the six-layer neocortex arose in mammals around 200 million years ago^1^, whereas distinct basal ganglia nuclei were already present in the last common ancestor of vertebrates more than 500 million years ago^2^.

Brain structures, circuits, and cell types have acquired adaptations and new functions along specific evolutionary lineages. Numerous examples of modifications to specific cell types within larger conserved brain systems have been discovered, including hindbrain circuits that control species-specific courtship calls in frogs^3^, the evolution of trichromatic vision in primates^4^, and neurons that have converted from motor to sensory processing to produce a novel swimming behavior in sand crabs^5^. Evolution can modify brain structures through a wide range of mechanisms, including increasing or reducing production of cells of a given type, altering the molecular and cellular properties of shared cell types, reallocating or redeploying cell types to new locations in the brain, or inventing entirely new cell types (Fig. 1a).

**Figure 1.**
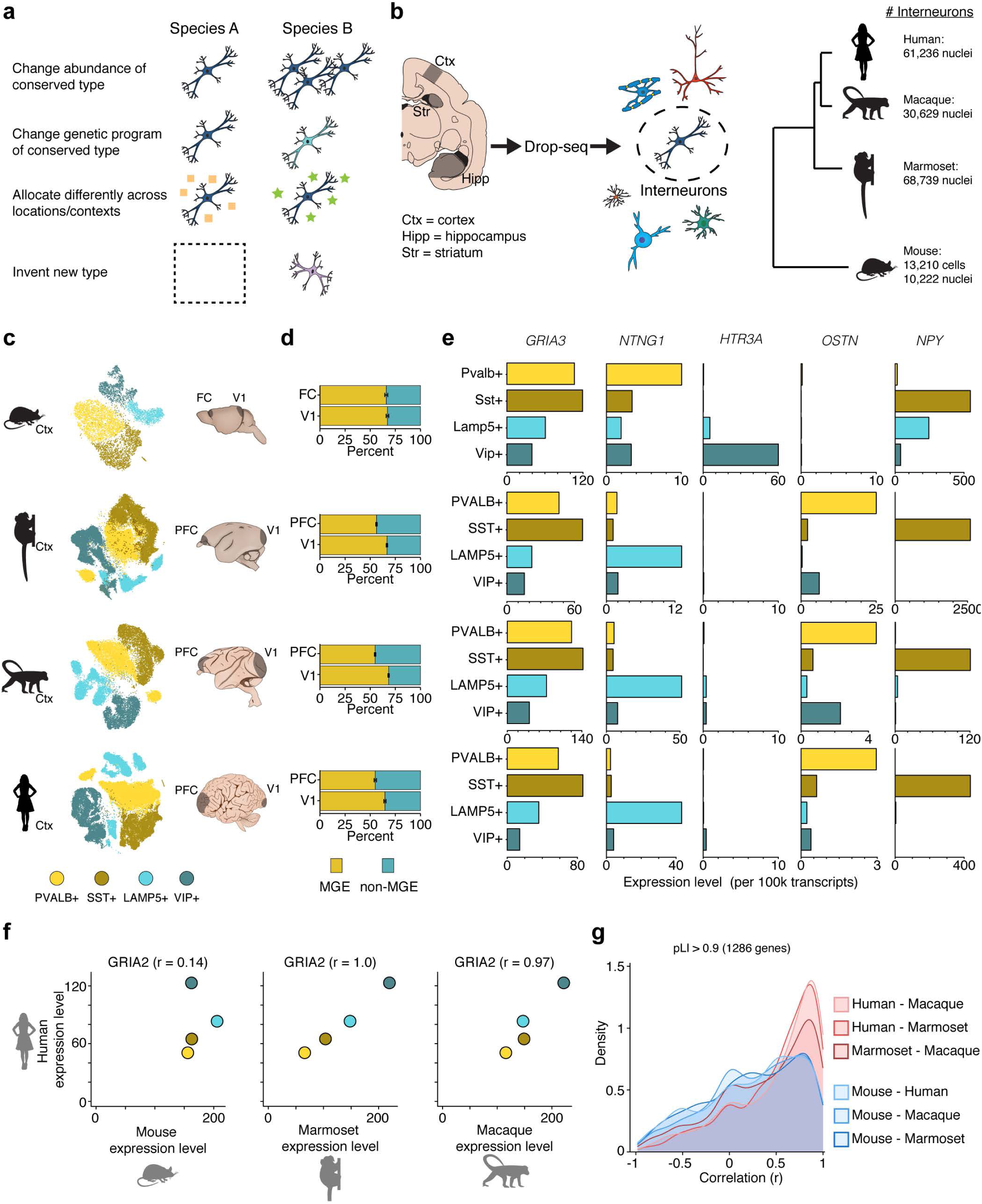
Analysis of cortical interneurons in mouse, marmoset, macaque, and human. **a**, Schematic showing possible modes of change in cellular assemblies across species. **b**, Schematic of experimental workflow and numbers of interneurons sampled in each species. Region abbreviations: Ctx = neocortex, Hipp = hippocampus, Str = striatum. **c**, t-distributed stochastic neighbor embedding (t-SNE) of cortical interneurons in each species. Cells are colored by membership in one of the four major neocortical classes: *SST+, PVALB+, VIP+*, or *LAMP5+* (dark brown or dark green cells represent the minority of cells that co-expressed *SST* and *PVALB* or *LAMP5* and *VIP*, respectively). **d**, The proportion of medial ganglionic eminence (MGE)-derived (*SST+* or *PVALB+*) and non-MGE–derived (*VIP+* or *LAMP5+*) types across two cortical regions (frontal/prefrontal cortex, PFC, and visual cortex, V1) for each species. Error bars represent 95% binomial confidence intervals. **e**, Examples of markers with different enrichment patterns across species (see also Extended Data Fig. 1b). Values are scaled expression levels (number of transcripts per 100k) for each of the four main cortical interneuron classes. **f**, Scaled expression levels (number of transcripts per 100k) for *GRIA2*, a gene encoding an AMPA receptor subunit, in human vs. mouse, marmoset, and macaque for the four major interneuron classes. Dots are colored as in (**e**). **g**, Density histograms showing correlation distribution of expressed genes between pairs of species (red = primate–primate pairs, blue = primate–mouse pairs). pLI = probability of loss of function intolerance.

Single-cell RNA sequencing, which systematically measures gene expression in thousands of individual cells, has recently enabled detailed comparisons of cell types and expression patterns between homologous brain structures separated by millions of years of evolution^4,6,7^ (non-single cell approaches have also yielded important insights in this domain, e.g. ^8^). For example, one recent study compared neocortical cells between humans and mice, identifying conserved and diverged features of many cell types^7^. It is not yet known which of the many differences between mouse and human brains are specific to humans, and which are shared among primates.

In this study, we compared interneurons, a major class of neurons present in all vertebrates, in mice and three primates: marmoset, macaque, and human, which span ∼90 million years of evolutionary divergence (Fig. 1b). Interneurons contribute to local circuit assemblies and provide the main source of inhibition in neuronal circuits by releasing the inhibitory neurotransmitter GABA. Interneurons are born subcortically from progenitors that reside in transient proliferative zones called the ganglionic eminences, including the medial and caudal ganglionic eminences (MGE and CGE), and migrate to the neocortex and to subcortical structures during development^9^. Interneurons are particularly interesting for comparative analysis because they are morphologically and physiologically diverse within any one species, but major types are shared across the amniotes^10^. In mice, the same interneuron types recur across functionally distinct neocortical regions^11,12^. An understanding of interneurons’ evolution in primates and rodents could guide the choice of models for studying how microcircuits and excitatory/inhibitory balance are affected in human brain disorders. Moreover, although the main developmental origins for interneurons appear conserved, we still do not know how interneurons are qualitatively and quantitatively allotted to their destinations, nor the extent to which local cues shape interneuron gene expression in different species.

### Identifying interneurons in mice and primates

We used Drop-seq^13^, a single-cell RNA sequencing technology, to measure RNA expression in nuclei of telencephalic brain cells (i.e., cells from brain regions including neocortex, hippocampus, and striatum) from adult animals of four species: mouse, common marmoset, rhesus macaque, and human. By applying unsupervised methods to the data from each individual species and from combinations of species, we classified transcriptionally distinct and similar groups of cells. We identified interneurons using canonical, conserved markers (e.g., *GAD1* and *GAD2*, which encode the glutamate decarboxylase required for synthesis of GABA) as well as class-specific molecular markers. In total, we sampled 68,739 telencephalic interneurons from marmoset, 61,236 from human, 30,629 from macaque, and 23,432 from mouse.

It was not previously known whether the broad classes of interneurons identified in mouse^11,12,14^ also exist in all three primates, and if so, whether they could be delineated with the same set of markers. Across all four species, the same four genes (*SST, PVALB, VIP*, and *LAMP5*) exhibited mutually exclusive expression while together accounting for almost 100% of neocortical interneurons, suggesting that these markers stably delineate a core repertoire of interneuron types among rodents and primates (Fig. 1c; see also^7^).

### Interneuron abundances and local specialization in neocortex

Within conserved brain structures, evolutionary changes in cell numbers or proportions can have profound functional consequences; such alterations appear to be major effectors of brain evolution^5,15,16^. Mammalian neocortex is patterned into functionally specialized fields, called areas, that differ in cytoarchitecture, cell number, and connectivity. There is a fundamental distinction between primary sensory areas of the neocortex, which process visual, auditory and tactile information as part of well-defined hierarchies, and association areas such as prefrontal cortex, which perform higher-order functions. Primates, and particularly humans, have neocortices that are disproportionately enlarged relative to those of other mammals^17^. The accompanying changes in cellular composition are not well understood, although recent quantitative stereological methods have begun to relate cortical specialization to cell-type composition across the neocortex^16^.

In primates, but not mice, frontal association areas (FC/PFC) differed from primary visual cortex (V1) in the extent to which interneurons were derived from MGE as opposed to other eminences (Fig. 1d). In all four species, V1 contained similar proportions of MGE (marked by *SST* or *PVALB*) and non-MGE interneurons (marked by *VIP* or *LAMP5)*: ∼66% MGE and ∼34% non-MGE. In primates, however, PFC harbored a significantly higher proportion of non-MGE interneurons (∼55% MGE, ∼45% non-MGE; Fig. 1c). Like PFC, association areas in temporal and parietal cortex contained proportionally more non–MGE-derived interneurons than V1 (Extended Data Fig. 1a). In primates, the upper neocortical layers have enlarged, particularly in association cortex^18^. Because MGE-derived interneurons preferentially populate deep layers^19^, the proportional increase in non–MGE-derived interneurons in primates is consistent with enlargement of upper-layer neocortical compartments and suggests greater recruitment of interneurons from the CGE to association cortex in primates.

### Genetic programs within conserved interneuron types

Homologous cell types can acquire species-specific functions through changes in gene expression^9,16,17^. To evaluate the extent to which gene-expression specializations distinguishing interneuron types are shared across species, we compared the expression level of each gene across the four principal interneuron classes (*PVALB*+, *SST*+, *LAMP5*+, *VIP*+) within each species, and then compared these class-specific relative expression profiles across the species surveyed. (Comparing within and then across species corrects for species-specific [e.g. sequence-related] influences on mRNA sampling, as well as for latent technical variables that might distinguish brains from different species.) This analysis enabled identification of conserved gene expression patterns. For example, *GRIA3*, which encodes an AMPA receptor subunit, was expressed in similar patterns across the four classes (*SST*+ > *PVALB+* > *ID2+* > *VIP*+) in all four species (Fig. 1e).

We first focused on genes that were selectively expressed at least one interneuron type (relative to the others) in at least one species (examples in Fig. 1e and Extended Data Fig. 1b). A clear pattern emerged: the great majority of human–mouse gene expression differences were shared among all three primates. For example, the neuropeptide Y (*NPY*) gene, a commonly used marker for specific interneurons, was expressed in both *SST+* and *LAMP5+* interneurons in mouse but was selectively expressed in *SST+* interneurons in marmoset, macaque, and humans. The netrin G1 (*NTNG1*) gene was, in mouse neocortex, selectively expressed in *PVALB+* interneurons; in primate neocortex, *NTNG1* was instead expressed by *LAMP5+* interneurons. In rare cases, a gene was enriched as a specific marker in one cell class in primates (e.g. *OSTN* in *PVALB*+ interneurons) but not detected at all in mouse interneurons^20^, or vice versa (e.g., *HTR3A*, which encodes serotonin receptor 3a, see also^7^). Other examples included synuclein gamma (*Sncg*), the short transient receptor potential channel 3 (*Trpc3*), and the IQ motif containing GTPase activating protein 2 (*Iqgap2*), which were expressed specifically in certain classes of interneurons in mice, but were either widely expressed (*SNCG*) or enriched in a different population (*TRPC3, IQGAP2*) among neocortical interneurons from primates (Extended Data Fig. 1b). Such cross-species expression variation has implications for choosing selective markers to define or characterize conserved cell types.

Far more genes were expressed in many or all neuronal types at quantitatively distinct levels. We know little about the extent to which the precise expression levels of such genes contribute to the specialized functions of neuronal types. However, understanding such relationships will be important for interpreting the significance of noncoding genetic variation in humans, as well as for selecting appropriate models for heterozygous mutations ascertained in human patients. A comparative lens can reveal the extent to which evolution has maintained a gene’s quantitative expression level in different cell types relative to one another. To evaluate conservation at this level, we identified 4051 expressed genes that exhibited at least 1.5-fold expression variation among the four main interneuron classes, and calculated the cross-species correlation of each gene’s expression measurements across those classes. Illustrating one of the main patterns revealed by this analysis, expression levels of *GRIA2*, a member of an AMPA receptor subunit family, exhibited relatively little variation (± 25%) among the four main interneuron classes in mice, but varied by 2–3-fold across the homologous interneuron types in primates, in a pattern (*VIP*+ > *ID2+* and *SST+* > *PVALB+)* that was highly correlated across the three primates (r = 0.95–1.0; Fig. 1f).

Genes that are dosage-sensitive in humans might have particularly strong evolutionary constraint on their expression levels. To evaluate this, we further focused on 1,286 genes that exhibit evidence of haploinsufficiency in humans as determined using pLI, a metric based on sequence variation across 60,706 human genomes that describes the probability that a given gene is intolerant of loss of function in human populations^21^. Relative expression of these human-haploinsufficient genes (pLI > 0.9) in interneuron types appeared to be much more constrained among primates than in the primate–mouse comparisons (Fig. 1g). (A control analysis revealed that the subset of genes with low pLI scores (pLI < 0.1) had correlation values around 0 for all species pairs, implying that genes that are tolerant of loss of function in humans are less constrained in their expression levels.) This relationship suggests that even dosage-sensitive genes have undergone substantial evolutionary change in their quantitative expression levels, and that these levels are more similar among primates than between primates and mice.

We also compared pairs of cell types in each pair of species, evaluating the extent to which differential-expression relationships were conserved between species across genes that are meaningfully expressed (> 10 transcripts per 100,000) in interneurons (Extended Data Fig. 2). Such comparisons offered abundant evidence that the relative expression levels (between interneuron subtypes) of a vast number of mutually expressed genes have been conserved. The overall correlation of relative expression levels between cell-type pairs was stronger for comparisons between primates than for comparisons between mice and any of the primates (Extended Data Fig. 2c).

### Regional specialization of expression patterns within the neocortex

To resolve types of interneurons at a finer scale, and to compare these types across species and brain regions, we used a computational approach, LIGER^22^, that aligns expression patterns across experiments and species (Extended Data Fig. 2), enabling us to compare gene expression programs within interneuron types across cortical regions (Fig. 2). In mouse, the expression programs of interneurons, in contrast to those of excitatory neurons, exhibit few differences across cortical locations^11,12,23^. In marmosets, each of 17 readily resolvable interneuron types was present in all seven cortical regions surveyed, confirming that, as in mouse, different cortical regions contain the same basic interneuron types (Fig. 2a). However, gene expression patterns for these conserved types differed across cortical regions. Across types, the median number of regionally differentially expressed genes (rDEGs, >3-fold difference) between PFC and V1 was 55 (Fig. 2b), exceeding the number observed when comparing mouse frontal and posterior cortical areas using the same criteria (median = 12.5, see also^11,12^).

**Figure 2.**
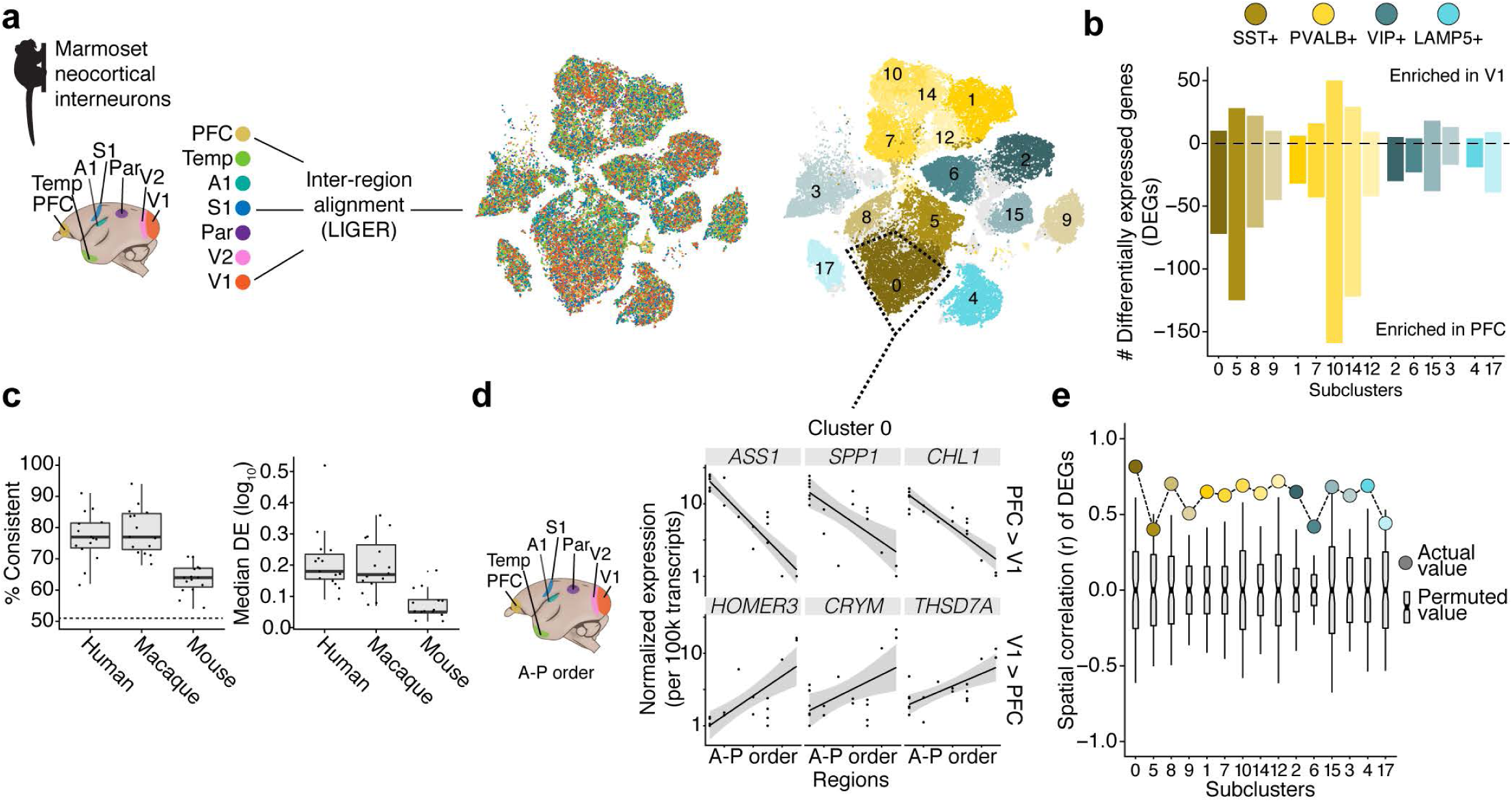
Comparing cortical interneurons within and across species. **a**, Integrative analysis of seven neocortical regions in marmoset using LIGER. t-SNE plots show resultant clusters with cells colored by cortical region of origin (left) and by cluster assignment (right). **b**, Histogram of the number of regionally differentially expressed genes (rDEGs) (>3-fold expression difference) between prefrontal (PFC) and primary visual cortex (V1) in each cell type (cluster) for which there were at least 50 cells per region. **c**, The rDEGs in marmoset (PFC, V1) tended to have the same regional differences in the other three species (left, percent of genes consistent with marmoset pattern; dashed line represents chance), with greater differential expression in humans and macaques than in mice (right). Dots represent rDEGs from each marmoset cluster; values were calculated from the cluster in the other species that had the most DE genes in common with that marmoset cluster. **d**, Normalized expression of rDEGs (identified between PFC and V1) across all seven neocortical regions in marmosets. X-axis arranged by anterior–posterior position of neocortical region. Plot shows the top three differentially expressed genes for the cluster outlined in **a** for each contrast (PFC>V1, V1>PFC). Dots are individual replicates within each region. **e**, Colored dots show averaged spatial correlations across rDEGs identified in each cluster when regions (n = 5, excluding PFC and V1) are arranged in anterior–posterior order. Gray boxplots show averaged correlations of the same rDEGs in each cluster when computed using permuted region orderings (n=120 possible orderings).

Although the number of rDEGs varied across clusters (Fig. 2b and Extended Data Fig. 3b), rDEGs identified for any one cluster or type tended to exhibit the same regional bias in the other clusters as well (Extended Data Fig. 3c,d). This suggests that most such differences reflect a common regional signature that is shared by diverse interneurons, rather than being specific to particular interneuron types. This regional bias did not extend to astrocytes (Extended Data Fig. 3e). The specific rDEGs defined in marmosets exhibited shared patterns of regional bias in interneurons in the other species: genes that were enriched in PFC vs V1 in marmoset were more likely to be more highly expressed in PFC than in V1 in the other three species, with greater probability and magnitude of difference in humans and macaques than in mice (Fig. 2c). These results suggest that interneurons acquire region-specific components of their molecular identities (see also^24,25^), and that these details are shared across species and most strongly among close relatives.

Spatial patterns of gene expression, including macro-scale gradients and the distinction between primary and higher-order neocortical areas, configure the layout of neocortical areas during development^26,27^ and persist into adulthood^28^. Gradients may contribute to excitatory neuron diversity^29^. Unlike excitatory neurons, which are born just below the neocortical areas in which they ultimately reside, neocortical interneurons are born subcortically and migrate into the neocortex post-mitotically; little is known about whether individual interneurons acquire information or specializations reflecting their ultimate areal locations within the neocortex. To explore this question, we investigated whether the expression of rDEGs identified in comparisons of PFC and V1 also varied across other neocortical regions. This analysis revealed a spatial logic: rDEG expression correlated strongly with anterior–posterior location (Fig. 2d); a control analysis in which region order was permuted yielded correlations distributed around zero (Fig. 2e). This anterior–posterior gradient is also correlated with, and might well reflect, the more complex patterns associated with the distinction between primary and higher-order neocortical areas^30,31^; for example, a number of genes had expression levels in parietal association cortex that were more similar to those in temporal and prefrontal cortex than those in the sensory areas more proximal to it. (Definitively parsing the overlapping effects of anterior-posterior, sensory-association, and other topographies would require a comprehensive sampling of neocortical areas.) Together, these results suggest that neocortical interneurons intrinsically detect and encode some aspect of their ultimate spatial position.

### Re-allocation of a shared interneuron type across brain structures

The main classes (*PVALB, SST, VIP, LAMP5*) of neocortical interneurons each contain many types^32^, and our analysis of the mouse and marmoset cortical interneurons (using LIGER) identified at least 15 transcriptionally distinct types (Extended Data Fig. 2b) with clear cross-species homologies in their global patterns of gene expression (Extended Data Fig. 4). These analyses affirm findings that homologous, molecularly-defined interneuron types can be identified across species spanning vast evolutionary distances, including reptiles, mice, and humans^7,33^. We have not attempted a definitive taxonomic classification here because we anticipate that improvements in single-cell technology and deeper ascertainment of neurons will further refine these categories.

The broad sharing of interneuron types, though, included notable differences. Notably, marmoset neocortex contained a substantial population of *LAMP5+* cells that co-expressed *LHX6* (Fig. 3a). The existence of these interneurons in the neocortex raised intriguing questions because *LHX6*, a transcription factor, participates in cell fate determination of MGE types, whereas *LAMP5+* neocortical interneurons come from the CGE^34^. In mouse, neocortical *Lamp5*+ interneurons consist of neurogliaform and single-bouquet types, which are the most numerous type of Layer 1 neuron and have distinct morphological, neurochemical, and connectivity properties^9^. Analysis by smFISH revealed that the spatial distribution of primate *LAMP5+/LHX6+* neurons was distinct from that of *LAMP5+/LHX6-* neurons, with the former tending to reside within the deep cortical layers (Fig. 3b). The proportion of interneurons that were *LAMP5+/LHX6*+ was 10-fold higher in marmoset, macaque, and human cortex than in mouse cortex, in all cortical regions analyzed (Fig. 3c). Thus, this cell type, recently reported to be much more abundant in human temporal lobe than in mouse primary visual cortex^7^, appears to have expanded throughout the neocortex in an ancestor of diverse primates.

**Figure 3.**
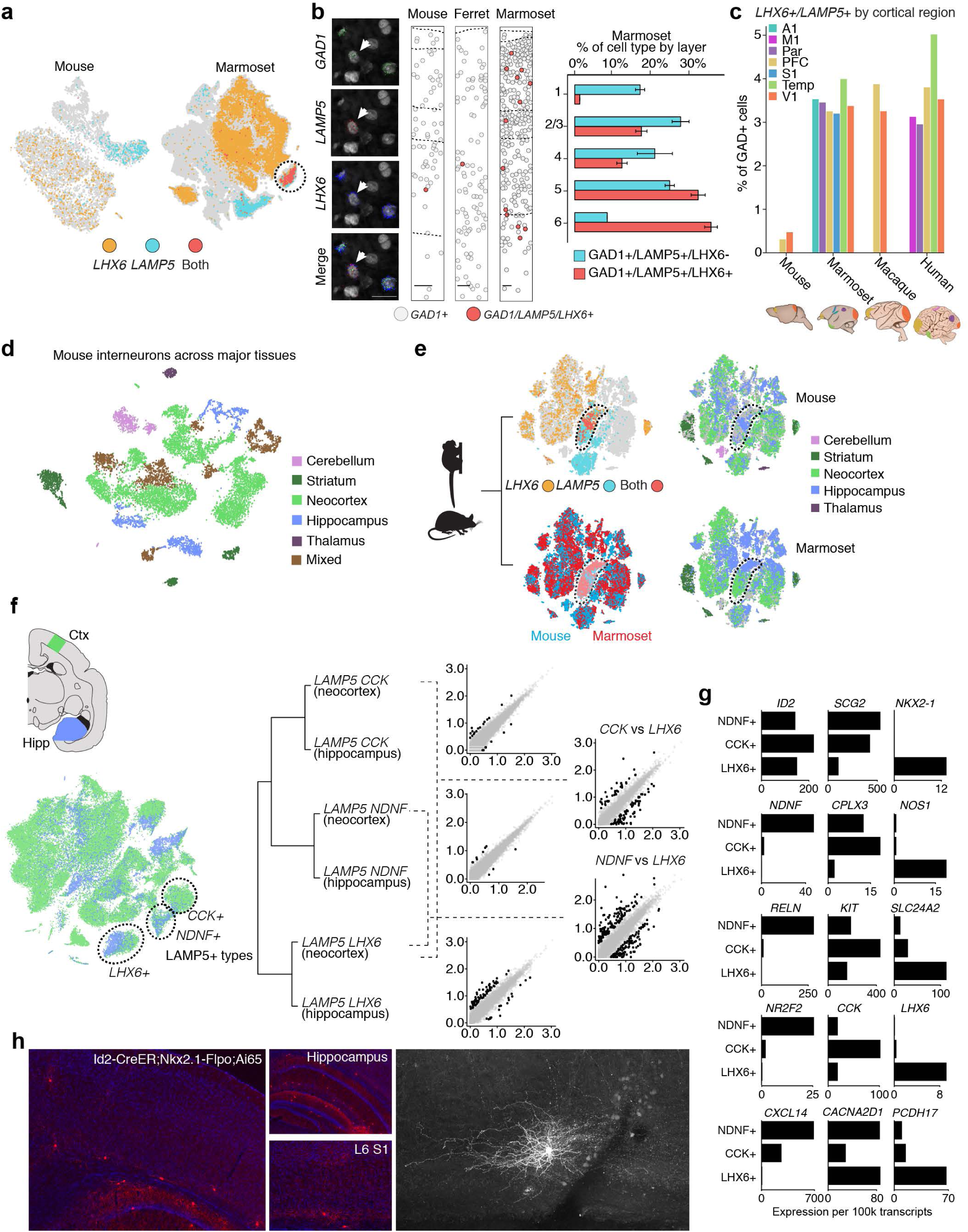
Cortical *LHX6+/LAMP5+* interneurons are more numerous in primates and are molecularly similar to conserved hippocampal interneurons. **a**, Cells expressing *LHX6, LAMP5*, or both are plotted on t-SNEs from mouse and marmoset neocortical data. **b**, (left) Single molecule fluorescence *in situ* hybridization (smFISH) in marmoset neocortex showing an example of an *LHX6+/LAMP5+/GAD1+* cell. (right) Quantification by layer of *LAMP5+/LHX6-/GAD1*+ cells (blue) and *LAMP5+/LHX6+/GAD1+* cells (red) in marmoset neocortex. **c**, Abundances of *LHX6+/LAMP5+* cells, expressed as proportions of *GAD1*+ interneurons sampled by Drop-seq, in marmoset, macaque, and human. Each sampled neocortical region is plotted separately. **d**, Mouse interneurons across major brain structures from ^12^. Clusters are colored based on their dominant region of origin. **e**, (left, top) Integrative cross-species analysis (using LIGER) of marmoset neocortical, hippocampal, and striatal interneurons and mouse interneurons from (**d**). (left, bottom) The same t-SNE, colored to show cells expressing *LHX6, LAMP5*, or both genes. Cluster outlined in t-SNE contains cells that express both *LAMP5* and *LHX6*. (right) The same t-SNE, colored for each species separately by cell region of origin. **f**, (left) Clustering of marmoset neocortex and hippocampal interneurons. (right) Hierarchical clustering of *LAMP5+* subtypes, separated by region (neocortex and hippocampus). Scatter plots of relative gene expression in pairs of subtypes (log_10_). **g**, Scaled, normalized expression of select gene markers that distinguish the three main *LAMP5+* types. **h**, To identify this population in mouse, and to determine whether the same Nkx2.1 lineage gives rise to such cells in hippocampus and neocortex, *Id2-CreER; Nkx2.1-Flpo; Ai65* animals were examined. (*left*) Overview including neocortex and hippocampus. (*middle, top*) Hippocampus was abundantly labeled. (*middle, bottom*) In neocortex, labeling was extremely sparse and mostly restricted to Layer 6. Labeled cells could be found rarely in L2/3, but not at all in L1. (*right*) A biocytin-filled mouse *Id2;Nkx2.1* interneuron in neocortical layer 6.

The *LAMP5/LHX6+* neurons could represent an innovation of primates or an ancestral condition lost by laboratory mice. The ferret, as a carnivore, serves as an outgroup relative to mice and primates. Analysis by smFISH for *LAMP5, LHX6*, and *GAD1* in ferret neocortex showed that, like mice, ferrets lacked a large deep layer *LAMP5+/LHX6*+ interneuron population, suggesting that the expansion of *LAMP5+/LHX6+* interneurons is a primate innovation (Fig. 3b).

To better appreciate the developmental and evolutionary origins of *LAMP5+/LHX6+* interneurons, we sought clues from other brain areas. Progenitors in the ganglionic eminences give rise to interneurons that migrate to the neocortex, striatum, hippocampus, and other subcortical structures. Comparing the expression profile of the primate cortical *LAMP5+/LHX6*+ population to expression profiles of 17,952 interneurons sampled from eight major structures of the mouse brain^12^ revealed that primate cortical *LAMP5+/LHX6+* cells most closely resembled *Lamp5+/Lhx6+* interneurons in the mouse hippocampus (Fig. 3d,e). Although neurogliaform interneurons in the neocortex are thought to derive solely from the CGE, mouse hippocampal *Lamp5+/Lhx6+* interneurons arise from the MGE; such neurons comprise the closely related ivy and neurogliaform subtypes, the most numerous hippocampal *Nos1*+ interneurons^35^.

To directly evaluate the similarity of the *LAMP5+/LHX6+* populations in hippocampus and neocortex, we analyzed marmoset neocortical and hippocampal interneurons together (Fig. 3f). The neocortical and hippocampal *LAMP5+/LHX6+* populations formed a common cluster, indicating that they were more similar to each other than to the other two *LAMP5*+ neocortical subtypes, which did not express *LHX6* (Fig. 3f,g). Notably marmoset hippocampal and neocortical *LAMP5+/LHX6*+ populations expressed *NKX2-1* (Fig. 3g), which in mouse is obligately downregulated in MGE-derived interneurons destined for the neocortex^36^ but persists in some human cortical interneurons^37^. Fate mapping of interneurons identified by *Id2/Nkx2-1* in mice confirmed that the cortical and hippocampal populations arose from a common (MGE) origin (Fig. 3h).

Primates might have evolved customized allocation for these *LAMP5+/LHX6+* cells from the MGE to neocortex specifically or could have simply expanded the generation of these cells for all brain structures, for example by expanding their progenitor pool. To evaluate these possibilities, we asked whether the *LAMP5+/LHX6+* population has also increased in primate hippocampus. smFISH analyses in in marmoset and mouse indicated that *LAMP5+/LHX6+* neurons populate the same hippocampal layers in the CA1/CA2 region (Extended Data Fig. 5, see also^38^). Thus, the ten-fold expansion of *LAMP5+/LHX6+* neurons in primates appears selective to the neocortex and is likely to represent differentially customized allocation (potentially via different rates or cues for migration, or via different rates of cell death) between neocortex and hippocampus relative to mice. Intriguingly, the *LAMP5+/LHX6*+ cells are distinct from, but molecularly most closely related to, recently described LAMP5+ cells that have acquired a distinct “rosehip” morphology and distinct physiological properties in human neocortex relative to their molecularly homologous cell population in mouse cortex^7,39^.

### A novel molecular interneuron type in primate striatum

Although the neocortex has greatly expanded and specialized in the primate lineage^18,40,41^, the basal ganglia are deeply conserved collections of subcortical nuclei – so much so that the lamprey, which shared a last common ancestor with mammals more than 500 million years ago, retains nuclei, circuitry, and basic cell types homologous to those observed in mice^2^. Therefore, we expected that interneuron types in the striatum (the largest part of the basal ganglia) would be highly conserved between primates and mice. To our surprise, marmoset striatum revealed, in addition to all the major classes of striatal interneurons found in mice^12,42^, a transcriptionally distinct type that expressed *VIP* and *TAC3.* This subtype did not appear to have a molecularly homologous population among mouse striatal interneurons (Fig. 4a-b). Because *VIP* is also sparsely expressed in other marmoset striatal interneuron types [Fig. 4b], we hereafter refer to the population as *TAC3+*. The *TAC3*+ interneuron population was present in male and female marmosets and in multiple striatal nuclei, including the caudate nucleus, putamen, and nucleus accumbens. Surprisingly, it constituted ∼30% of all interneurons in the striatum.

**Figure 4.**
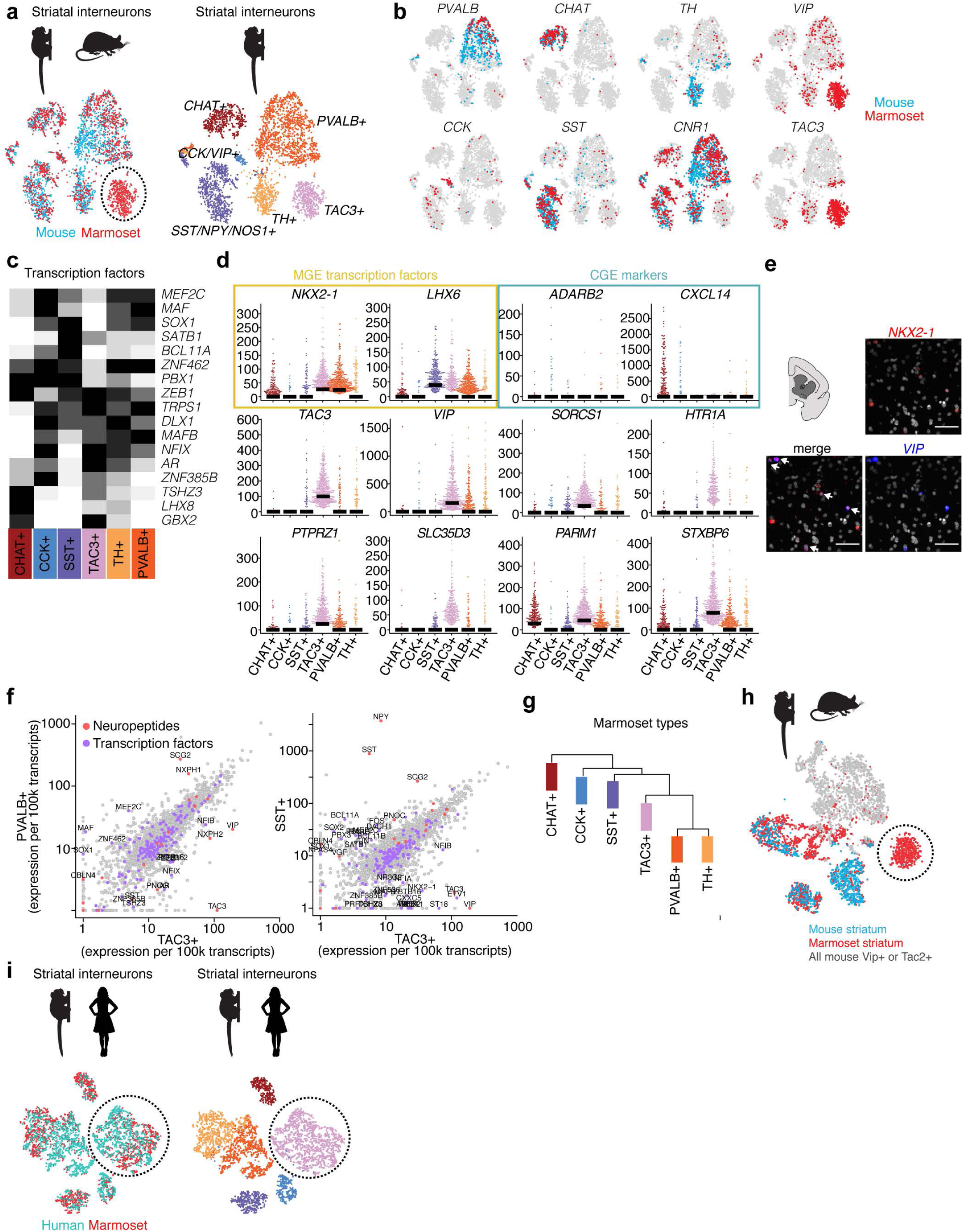
A primate striatal interneuron type not observed in mouse. **a**, Integrative cross-species alignment (using LIGER) of marmoset and mouse striatal interneurons. **b**, Cells expressing markers for each interneuron cluster plotted for marmoset (red) and mouse (blue). **c**, Heat map of expressed transcription factors in marmoset for each striatal subtype. Each gene is scaled to its maximum value across types (black = max). **d**, Beeswarm plots showing additional markers that distinguish *TAC3+* interneurons from other interneuron types in marmoset, including MGE transcription factors (shaded yellow) and CGE markers (shaded blue). Dots are individual cells; bars indicate median expression. **e**, smFISH for *VIP* and *NKX2-1* in marmoset striatum. Cells that co-express both genes identified by arrows. **f**, Scatterplots showing gene expression (log_10_) between *TAC3+* and *PVALB+* or *TAC3+* and *SST+* populations. Differentially expressed (> 3-fold difference) neuropeptides and transcription factors are labeled. **g**, Hierarchical clustering of all expressed genes within marmoset striatal subtypes. **h**, The analysis in (**a**) repeated, but additionally including all mouse extra-striatal interneurons from ^12^. For display, the t-SNE shows marmoset striatal interneurons (red), mouse striatal interneurons (blue), and any extra-striatal mouse interneuron that expressed *Vip* or *Tac2* in the Saunders et al. ^12^ dataset (gray). Circled cells indicate marmoset *TAC3+* population. **i**, LIGER analysis pooling marmoset striatal interneurons with caudate interneurons isolated from human postmortem donors (n=2). Circled cluster indicates aligned marmoset and human *TAC3+* populations.

*TAC3+* interneurons expressed unique combinations of transcription factors, neuropeptides, transporters, and receptors that were not observed in other interneuron subtypes in marmosets (or in any subtype in mice) (Fig. 4c–f). These included genes encoding the androgen receptor (*AR*), serotonin receptor 1A (*HTR1A*), and sugar transporter *SLC35D3*.

The genes expressed by this novel population of striatal interneurons could provide hints about their developmental origins. Although *Vip* and *Tac2* (the mouse homologue of *TAC3*) are neuropeptide genes associated with CGE origin in mouse^9^, the marmoset *TAC3+* population expressed the MGE-associated transcription factors *LHX6* and *NKX2-1* (Fig. 4d). Because transcription factor and neuropeptide expression offered divergent clues, we used the entire genome-wide expression pattern to identify the interneurons most similar to the *TAC3+* interneurons. Hierarchical clustering situated the *TAC3+* population between the *SST+* and the *TH+* and *PVALB*+ populations, all of which are MGE-derived (Fig. 4g). This suggests that despite expressing some CGE-associated neuropeptides, the *TAC3+* cells are more similar to striatal MGE-derived types than CGE-derived types.

To evaluate the possibility that a population of cells homologous to the *TAC3*+ marmoset striatal interneurons might reside elsewhere in the mouse brain, as in the case of *LAMP5+/LHX6+* neocortical interneurons, we jointly analyzed the RNA expression profiles of marmoset striatal interneurons and all interneurons from eight regions of the mouse brain (from ^12^). The marmoset *TAC3+* population still formed its own cluster, suggesting that no homologous cell population existed in any of the mouse brain regions sampled (Fig. 4h). We also confirmed, by single-nucleus RNA sequencing, that no such population is present in ferret striatum, consistent with the interpretation that this cell population was introduced in the lineage leading to primates rather than being lost in mice.

The *TAC3*+ interneuron population appeared to be shared between marmosets and humans: comparing marmoset striatal interneurons to interneurons obtained from the caudate nucleus from adult human postmortem donors identified a human striatal interneuron population with a homologous pattern of gene expression (Fig. 4i and Extended Data Fig. 6). The genes that were differentially expressed in marmoset between the *TAC3*+ population and other striatal interneuron types also tended to be differentially expressed in the corresponding comparisons in human (Extended Data Fig. 6c). The *TAC3*+ population constituted 38% of the interneurons sampled in human striatum.

The abundance of the novel, TAC3+ interneuron population raised the question of whether it had replaced, or added to, conserved interneuron populations. The primates exhibited expanded representation of interneurons in the striatum overall: while interneurons were 4.1% of all striatal neurons in mice, they were 13.1% of all striatal neurons in marmosets and 10.8% in humans, consistent with stereological estimates of higher interneuron proportions in primate striatum^43^. Thus, compared to mice, primate striatum has more than doubled the proportion of interneurons, including an interneuron type with no clear homolog in mice.

## DISCUSSION

“Cell types” have been defined as collections of cells that change together over the course evolution^44,45^. In this study, we found that although most of the major molecularly defined types of cortical interneurons are conserved across mice, humans, marmosets, and macaques, these interneurons nonetheless have undergone surprising levels of evolutionary change in the genes they express and the relative levels at which they express pan-neuronal genes. The significance of these changes for the detailed physiological and connectivity properties of interneurons will be important to understand^46^.

We found that the primate striatum contains an abundant interneuron type that has no homologous cell population in mice. These TAC3+ interneurons constituted 30% of interneurons in human and marmoset striatum, and expressed a suite of transcription factors and neuropeptides that distinguished them from other striatal interneurons. This innovation in primate striatum was accompanied by a broader expansion in the numbers of interneurons that doubled their representation as a fraction of all striatal neurons. These observations raise questions about how expanded numbers and types of interneurons have changed the functions of striatal microcircuits and their roles within the larger cortico-striatal circuits that contribute to primate cognition and potentially to neuropsychiatric disorders^47^.

We found that an interneuron type that is abundant in the mouse hippocampus—the ivy cell, which has properties similar to neurogliaform cells and is defined by co-expression of *Lamp5* and *Lhx6*^35^—has expanded throughout the neocortex in primates. Primates have retained ivy cells in the hippocampus but appear to have also greatly upregulated the production of these cells, increased their recruitment to the neocortex, and expanded their distribution throughout neocortical areas and layers. Other neurogliaform cell types in mouse neocortex are preferentially found in upper layers and signal by volume transmission, the diffuse release of the inhibitory neurotransmitter GABA in the absence of conventional synapses^48^. Given that these cells were proportionally most numerous in the deep layers of the primate neocortex, one possibility is that in primates, these cells now contribute diffuse inhibitory signaling in new neocortical contexts.

The qualitative and quantitative deployment of gene expression across conserved interneuron types indicated that, even for genes that are pan-neuronally expressed, evolution has strongly constrained quantitative gene expression levels on evolutionary time scales, although substantial differences in gene expression and cellular proportions still clearly distinguish primates from mice.

Efforts to model the effects of specific genes and mutations on human brain function and illness could be facilitated by systematic data sets that reveal the extent to which each gene’s cell-type– specific pattern of expression is shared by humans with each other species (e.g., ^8^). We hope that the data from the experiments reported here will inform the design and interpretation of such studies. Accordingly, we have developed a simple, web-based data resource to enable cross-species comparisons of interneurons (http://interneuron.mccarrolllab.org/).

Our results reveal the ways in which the cellular and molecular repertoires of mouse and primate neurons have adapted over time. These evolutionary paths likely involved diverse developmental mechanisms, including alteration of neurogenesis rates, the creation of novel migratory pathways, and changes in gene regulation. Their effects on circuitry, cytoarchitecture, and physiology will be important and interesting to understand.

The innovations among interneurons are notable because the single-cell expression studies of tetrapod species performed to date – in lizards, turtles, mice, and primates – had suggested that the known interneuron types are conserved across a broad taxonomic range. In this study, however, we identified surprising variation in interneurons within and across species, which was discordant with expectations in key ways. For example, the primate CGE might be expected to harbor evolutionary novelties in its interneuron repertoire because it generates a larger proportion of interneurons in primates than in rodents^49^, because CGE interneurons are born later than MGE interneurons, and because CGE interneurons preferentially occupy the expanded upper neocortical layers^50^. However, the most striking cellular aspects of rodent–primate divergence in the cortex, hippocampus, and striatum involved interneurons whose RNA expression patterns indicated that they originated in the MGE. Similarly, although the neocortex has attracted intense interest because it is a highly evolved and specialized structure that is thought to underlie expansions in primate cognitive capability, it was in the striatum that we identified a primate interneuron type with no mouse homolog. The systematic analysis of many more species and cell types may reveal more such examples of evolutionary flexibility and innovation.

## Acknowledgments

This work was supported by the Broad Institute’s Stanley Center for Psychiatric Research and Brain Initiative grant U01MH114819 to G. Feng and S.A.M, by the Dean’s Innovation Award (Harvard Medical School) to G. Fishell and S.A.M., and by the Hock E. Tan and K. Lisa Yang Center for Autism Research at MIT, the Poitras Center for Psychiatric Disorders Research at MIT and the McGovern Institute for Brain Research at MIT (G. Feng). Also supported by NINDS RO1NS032457 (C.A.W.). C.A.W. is an Investigator of the Howard Hughes Medical Institute. We thank Dr. Maude W. Baldwin, Avery D. Bell, Steven Burger, Dr. Christopher Patil, and Dr. Randy L. Buckner for comments on manuscript drafts; Dr. Christian Mayer for analysis advice; and Dr. Christina Usher for assistance with manuscript preparation.

## Author Contributions

F.M.K, S.A.M., G. Feng, and G. Fishell designed the study. F.M.K. prepared and dissected tissue; L.B. and M.G. developed the nuclei Drop-seq protocol. M.G., A.L., C.D.M., N.R., E.B., and L.B. performed Drop-seq and prepared sequencing libraries. M.G. performed sequencing, alignment, and QC analysis. F.M.K., A.S., J.N., A.W., D.K. R.d.R., and S.A.M. developed analysis pipelines. F.M.K. analyzed the data with input from S.A.M, G. Fishell, M.F., A.L. and A.S. D.K. developed the web resource. Q.Z., C.W., M.B., V.T., R.S., C.A.W., L.K., S.B., and G. Feng provided tissue for Drop-seq and smFISH experiments. K.L., H.Z., C.D.K., N.R., E.B., M.F-O., J.L., F.M.K. and J.D. performed and analyzed smFISH experiments. R.M., B.S., and B.R. contributed fate-mapping experiments. F.M.K and S.A.M wrote the paper with input from coauthors.

---

**Extended Data Table 1.** Specimen information table.

**Extended Data Table 2.** Reagent and resource table.

**Extended Data Figure 1.**
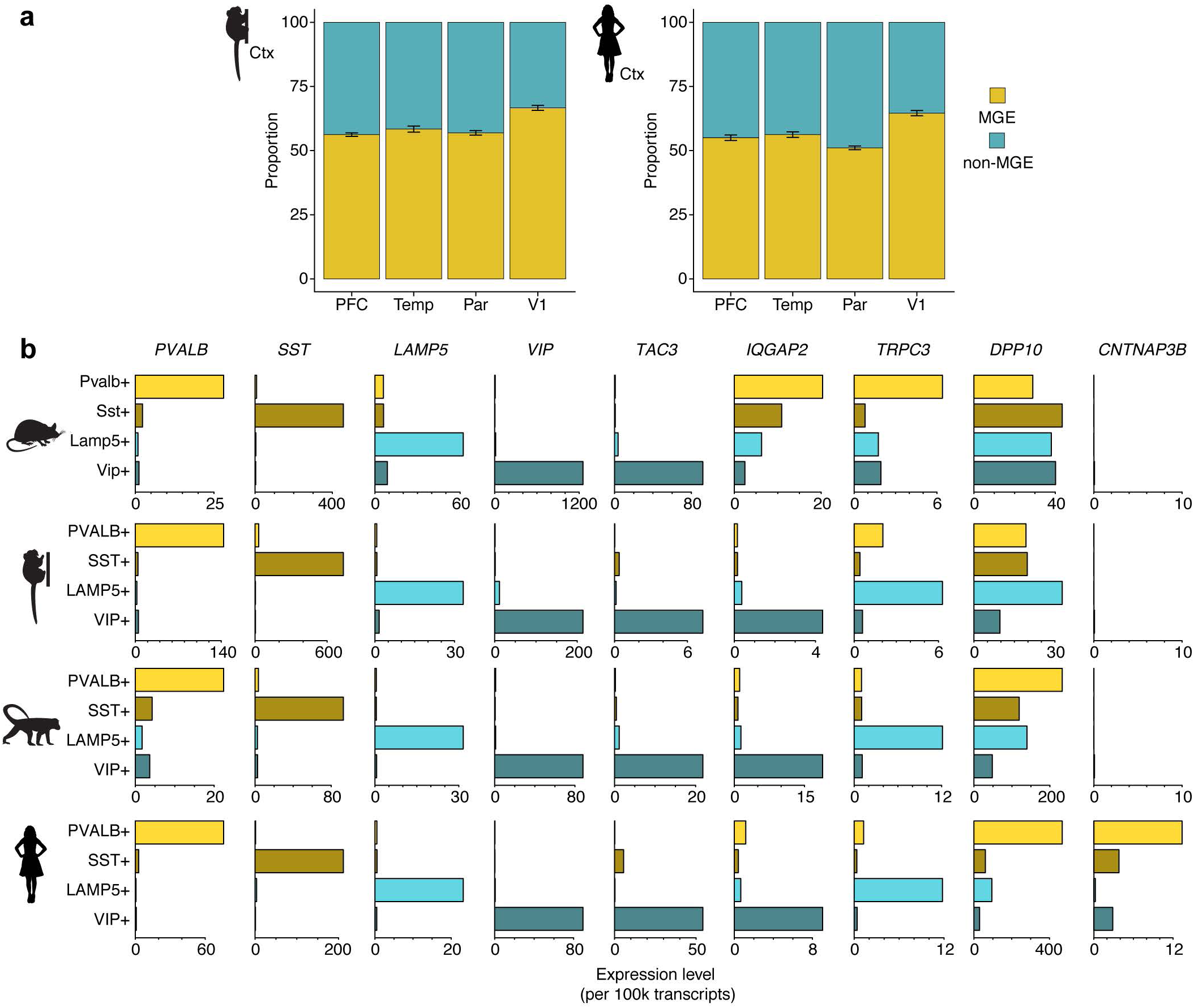
Interneuron abundances and gene expression in neocortex. **a**, Proportion of MGE and non-MGE interneurons in cortical association areas (prefrontal cortex, temporal pole, and lateral parietal association cortex) and in primary visual area (V1) in marmoset and human. Error bars represent binomial confidence intervals. **b**, Examples of markers that are consistent, or that vary across species, within the four primary interneuron classes. Values are scaled counts per 100k transcripts.

**Extended Data Figure 2.**
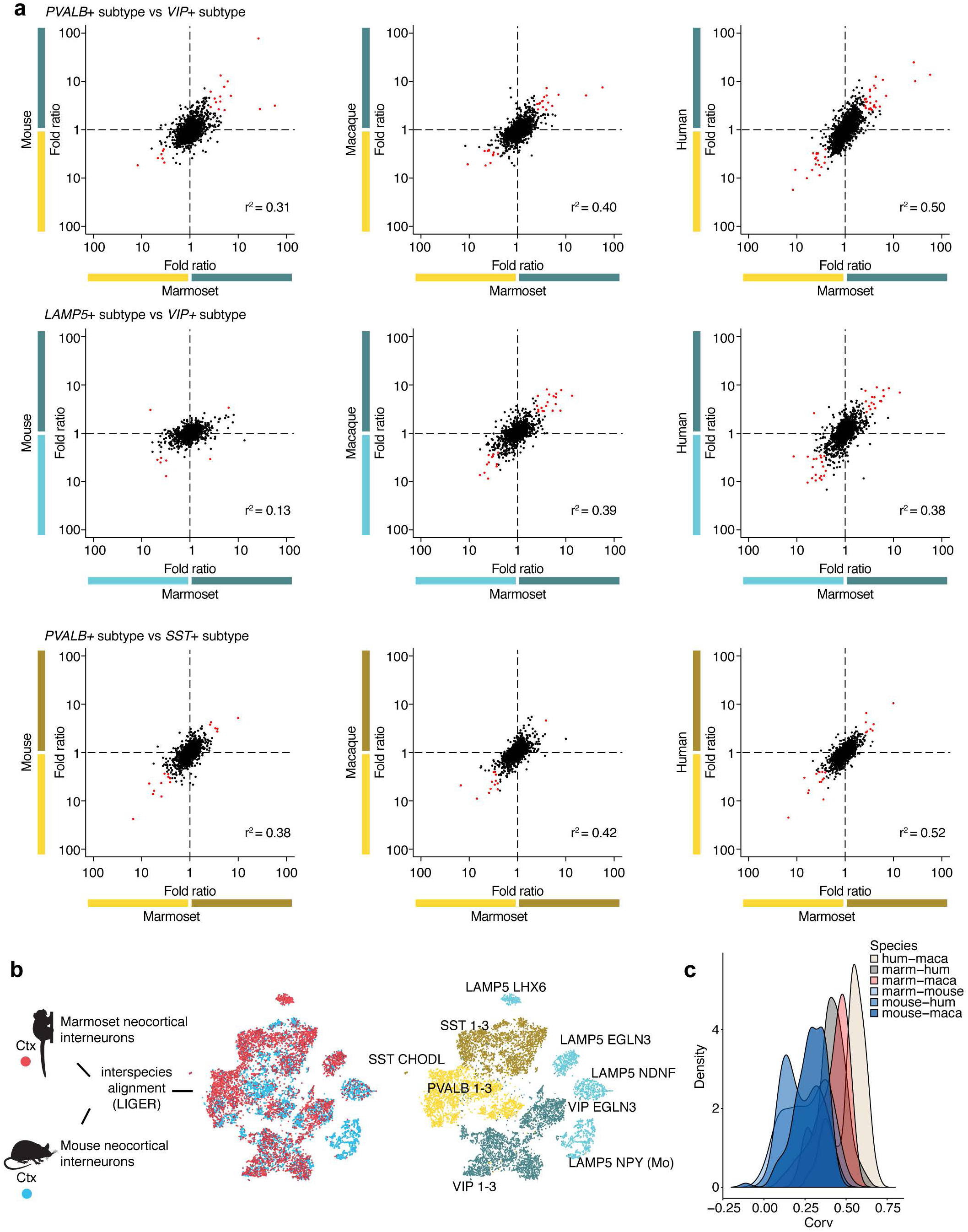
Pairwise comparisons of neocortical interneuron types across species. **a**, Fold difference of each expressed gene between types of MGE-derived interneurons and types of CGE-derived interneuron across pairs of species. Genes in red have >3-fold expression difference in either cell type in each species pair. **b**, (left) LIGER integration of marmoset (red dots) and mouse (teal dots) neocortical interneurons. (right) Same t-SNE with clusters colored by interneuron class (colors as in Fig. 1). **c**, Density histogram of correlation (*r*) values of fold differences computed for each possible cluster pair from LIGER species-integrated analyses. Each density trace corresponds to a species pair; blue traces indicate primate–mouse comparisons, and other colors indicate primate–primate comparisons.

**Extended Data Figure 3.**
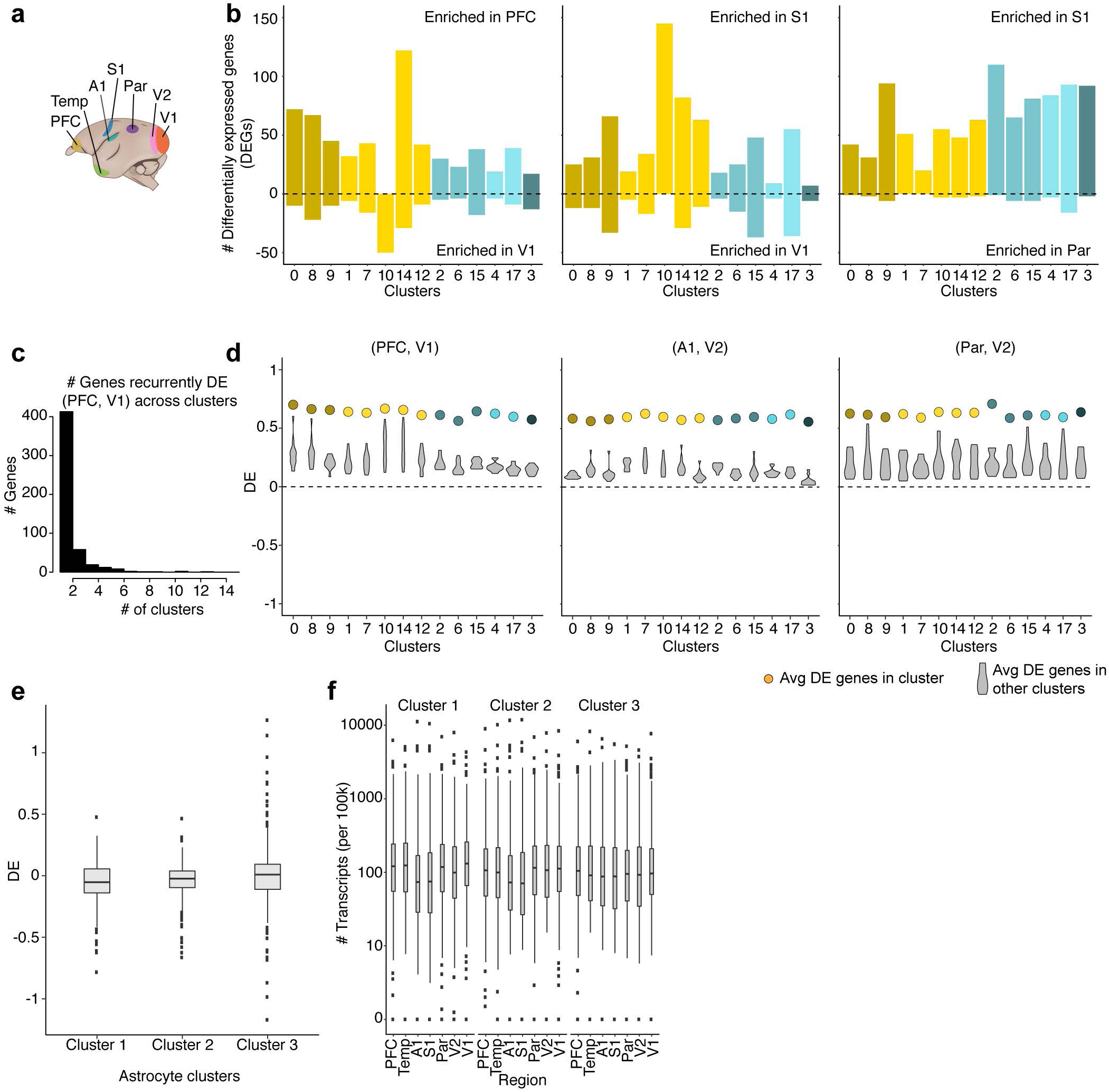
Regional gene expression variation in neocortex. **a**, Schematic of neocortical region locations in marmoset. **b**, Histogram of the number of regionally differentially expressed genes (rDEGs) (>3-fold expression difference) between three representative pairs of regions, in each cell type (cluster) for which there were at least 50 cells per region. **c**, Histogram of the number of interneuron clusters (cell types) in which a given gene is differentially expressed. At a threshold of >3-fold, most genes are only differentially expressed in a single cell type (cluster). **d**, Colored dots represent average fold difference of DEGs in each cluster in marmoset interneurons. Violin plots represent the distribution of average fold differences in each cluster (cell type) when using rDEGs from other clusters. Three representative region pairs are shown. **d**, Fold ratios (log_10_) between PFC and V1 for three astrocyte subtypes (n = 32,600 nuclei) in marmosets using rDEGs identified between PFC and V1 in interneurons. **e**, Expression in astrocytes of genes that exhibited an anterior–posterior expression gradient in interneurons. Regions are arranged in anterior–posterior order on the x axis.

**Extended Data Figure 4.**
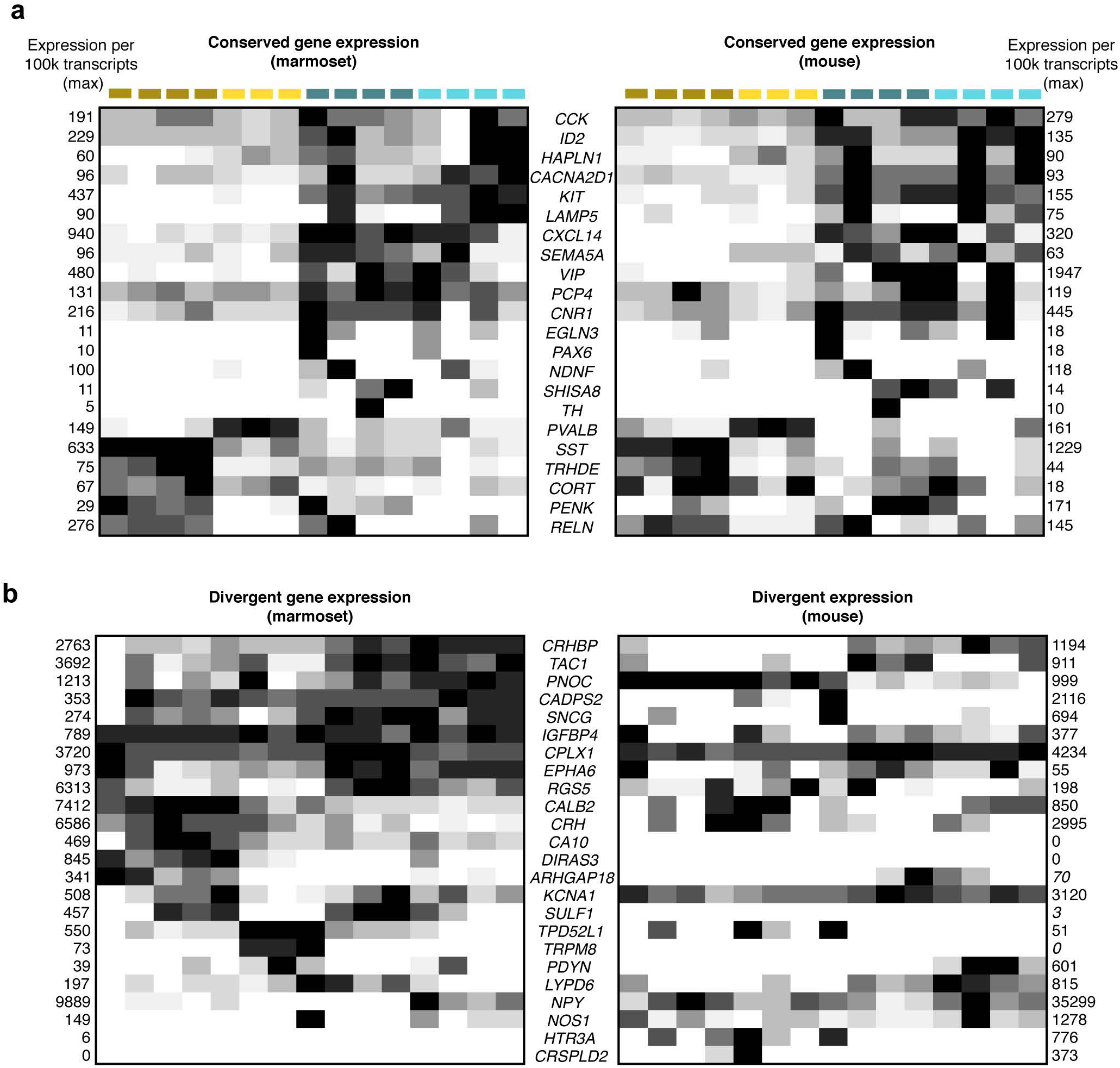
Conserved and divergent gene expression across neocortical types. **a**, Heat map of exemplar genes that had consistent patterns of expression in LIGER integrated marmoset–mouse clusters from Extended Data Fig. 2b. Each gene (row) is scaled to the scaled max (black) expression (values given outside plots) for each species separately. **b**, Heatmap of exemplar genes that have divergent expression patterns in LIGER-integrated marmoset–mouse clusters from Extended Data Fig. 2b. Each gene (row) is scaled to the scaled max (black) expression (values given outside plots) for each species separately.

**Extended Data Figure 5.**
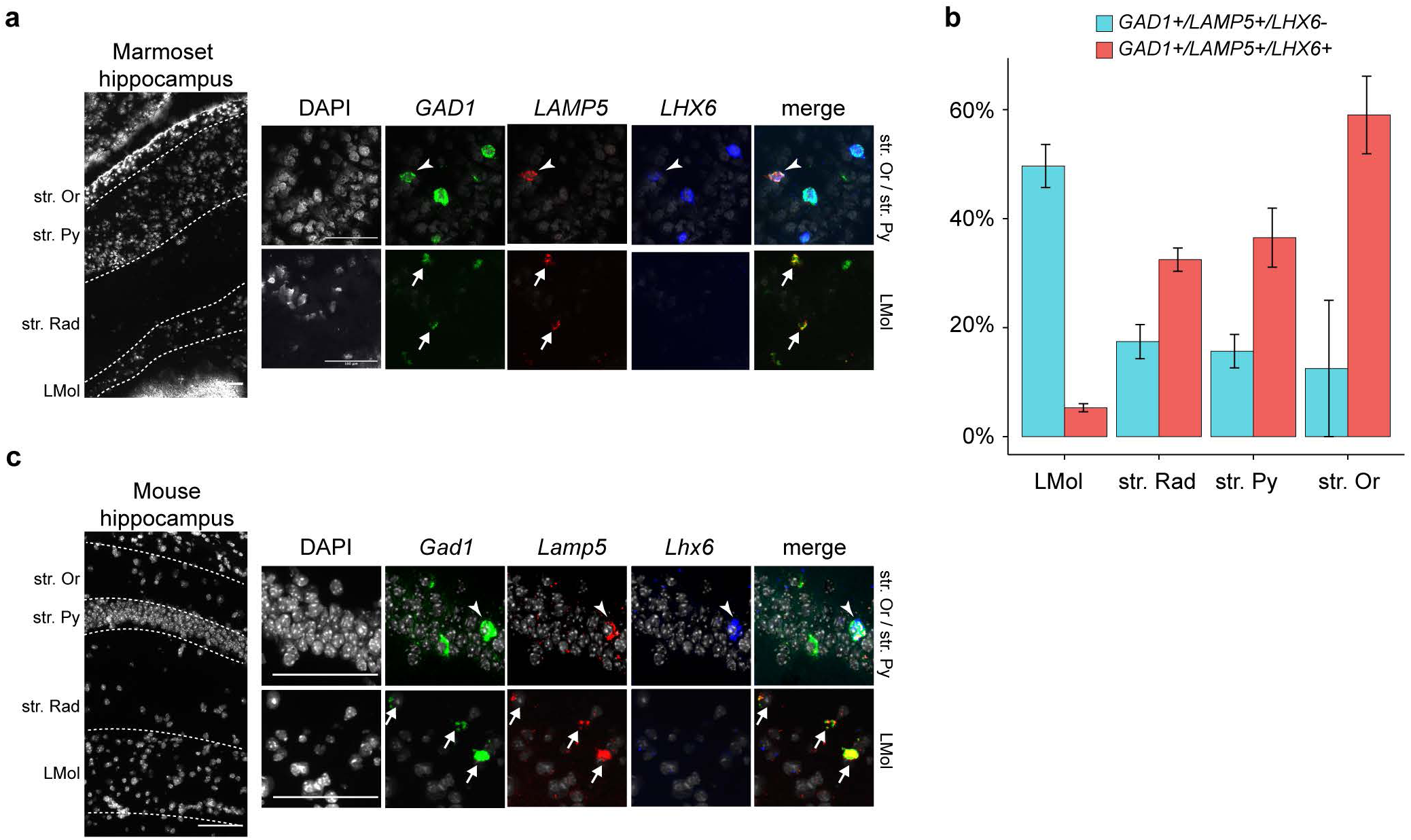
LAMP5+ interneuron types in hippocampus. **a**, Single-molecule fluorescence in situ hybridization (smFISH) for *GAD1, LAMP5*, and *LHX6* in marmoset hippocampal layers (CA1/CA2 subfields). Arrowhead indicates triple positive cells; arrow indicates the *LHX6*-population. (*top row)* Strata oriens (Str. Or) and strata pyramidale (Str. Py). (*bottom row)* strata lacunosum moleculare (LMol). Scale bars = 100 um. **b**, Quantification of *GAD1*/*LAMP5/LHX6+* (green) and *GAD1*/*LAMP5/LHX6-* cells as percentage of all *GAD1+* cells in marmoset hippocampus (compare to mouse data in^38^). **c**, smFISH for *Gad1, Lamp5*, and *Lhx6* in mouse hippocampal layers (CA1).

**Extended Data Figure 6.**
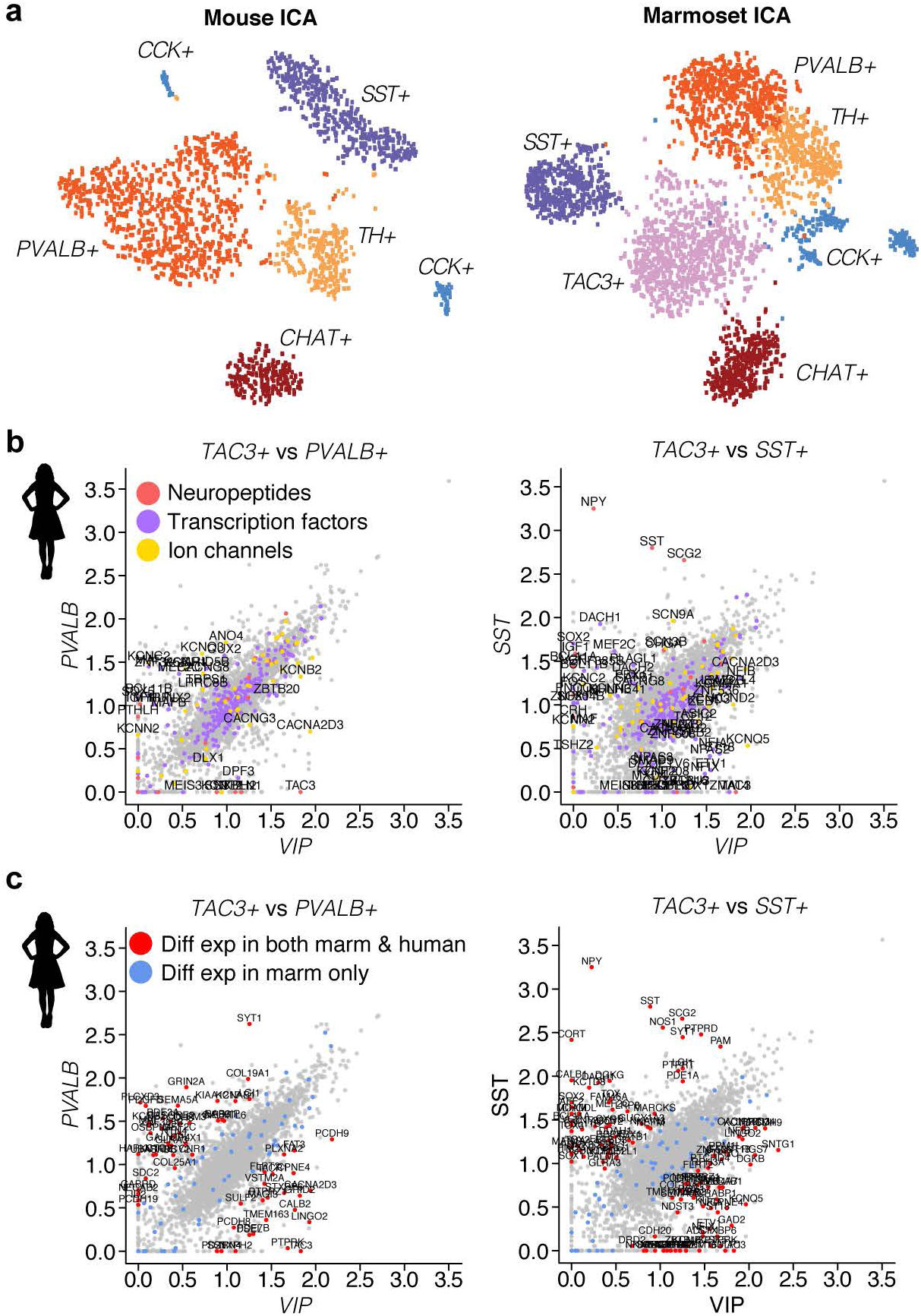
Interneuron types in striatum. **a**, t-SNE representations of ICA-based clustering for mouse and marmoset striatal interneurons. **b**, Gene expression differences in human caudate between *TAC3*+ and *PVALB*+ (left) or *TAC3*+ and *SST*+ (right) populations. Neuropeptides (red) and transcription factors (blue) are labeled. **c**, Same data as in **b**, but instead highlighting genes that were differentially expressed in both marmoset and human (red), or only in marmoset (blue).

## METHODS

Specimen information is available in Extended Data Table 1.

Reagent information is available in Extended Data Table 2.

### Specimens and Donors for Nuclei Drop-seq

#### Mouse

Mouse experiments were approved by and in accordance with Harvard Medical School IACUC protocol number IS00000055-3. Sections of frontal and visual cortex were prepared from male and female adult mice (60–70 days old; C57Blk6/N, Charles River Labs Stock #027). Mice were deeply sedated with isoflurane and transcardially perfused with ice-cold Sucrose-HEPES buffer described in^1^, which contains (in mM) 110 NaCl, 2.5 KCl, 10 HEPES, 7.5 MgCl_2_, 25 glucose, 75 sucrose (∼350 mOsm/kg^-1^); sectioned; and flash-frozen on liquid nitrogen.

#### Marmoset

Marmoset experiments were approved by and in accordance with Massachusetts Institute of Technology IACUC protocol number 051705020. Three adult marmosets (1.5–2 years old; one male, 2 females) were deeply sedated by intramuscular injection of ketamine (20-40 mg/kg) or alfaxalone (5-10 mg/kg), followed by intravenous injection of sodium pentobarbital (10– 30 mg/kg). When pedal withdrawal reflex was eliminated and/or respiratory rate was diminished, animals were transcardially perfused with ice-cold Sucrose-HEPES buffer. Whole brains were rapidly extracted into fresh buffer on ice. Sixteen 2-mm coronal blocking cuts were rapidly made using a custom-designed marmoset brain matrix. Slabs were transferred to a dish with ice-cold Dissection Buffer^1^, and regions of interest were dissected using a marmoset atlas as reference^2^. Regions were snap-frozen in liquid nitrogen and stored in individual microcentrifuge tubes at - 80°C.

#### Macaque

Whole brains from two healthy, immunologically and treatment-naive adult macaques (2 males; 10–11 years old) were obtained from terminal experiments (IACUC 4315-02). Animals were deeply sedated with ketamine and euthanized by pentobarbital overdose, and transcardially perfused with ice-cold Sucrose-HEPES buffer. Brains were rapidly blocked in ∼5-mm coronal slabs and frozen in liquid nitrogen or isopentane on dry ice.

#### Human

Frozen tissue was obtained from the Harvard Brain Tissue Resource Center (HBTRC; McLean Hospital). Four donors were used for analysis of striatal interneurons, and two for analysis of neocortical interneurons. History of psychiatric or neurological disorders was ruled out by consensus diagnosis carried out by retrospective review of medical records and extensive questionnaires concerning social and medical history provided by family members. Several regions from each brain were examined by a neuropathologist. The cohort used for this study did not include subjects with evidence of gross and/or macroscopic brain changes, or clinical history, consistent with cerebrovascular accident or other neurological disorders. Subjects with Braak stages III or higher (modified Bielchowsky stain) were not included. None of the subjects had significant history of substance dependence within 10 or more years of death, as further corroborated by negative toxicology reports.

### Nuclei Drop-seq library preparation and sequencing

Nuclei suspensions were prepared from frozen tissue and used for Nuclei Drop-seq following the protocol described at https://protocols.io/view/extraction-of-nuclei-from-brain-tissue-2srged6. Drop-seq libraries were prepared as previously described^3^ with modifications, quantification, and QC as described in^1^, as well as the following modifications optimized for nuclei: in the Drop-seq lysis buffer, 8 M guanidine hydrochloride (pH 8.5) was substituted for water, nuclei were loaded into the syringe at a concentration of 176 nuclei/µL, and cDNA amplification was performed using around 6000 beads per reaction (15 PCR cycles were used for marmoset nuclei, and 16 for macaque and human nuclei). Raw sequencing reads were aligned to the following genome assemblies: GRCm38.81 (mouse), calJac3 (marmoset), Mmul8.0.1 (macaque), and hg19 (human). Reads that mapped to exons or introns of each assembly were assigned to annotated genes.

### Mouse single-cell dataset

Interneurons were curated *in silico* from the single-cell datasets available in ^1^ from available structures: frontal and posterior neocortex, striatum, cerebellum, thalamus, hippocampus, substantia nigra, and entopeduncular nucleus.

### Single species independent component analysis (ICA)

Initial analyses to identify interneurons based on marker expression were conducted on each species separately. Nuclei with fewer than 300 detected genes were removed from analysis. Briefly, independent component analysis (ICA, using the fastICA package in R) was performed on each species and each region’s digital gene expression (DGE) matrix separately after normalization and variable gene selection as in ^1^. These first-round individual-species analyses produced clustering solutions with ∼8–11 clusters of major cell types (neurons, glia, vasculature), from which interneuron clusters could be identified based on canonical markers (e.g. *GAD1, GAD2*). The raw DGEs were subsetted to include only cells from these clusters to form new, interneuron-only DGEs. Normalization, variable gene selection, and ICA was repeated on these interneuron-only DGEs, but this time the full ICA curation pipeline described in ^1^ was used to identify doublets, outliers, artifactual signals, and biological components of interest. Cells identified by this procedure as doublets or outliers were removed from the DGEs, and these filtered DGEs were then carried forward for integrated analyses across regions and/or species using LIGER^4^.

### Interneuron abundances and local specialization in neocortex

Proportions of *PVALB+* or *SST+* (MGE-derived) and *VIP+* or *LAMP5+* (non-MGE derived) were calculated for each species separately for frontal association areas (FC, mouse) or prefrontal cortex (PFC, primates) and visual cortex (V1). Cells were allocated to MGE or non-MGE pools based on their cluster assignment in individual-species ICA clustering. Error bars represent 95% confidence intervals for binomial probability, computed with the R package Hmisc.

### Identification of conserved and divergent genetic programs within conserved interneuron types

Interneurons from the neocortex of each species were partitioned into four main classes based on marker expression (*VIP, LAMP5, PVALB, SST*). (We also detected a rare and distinct, fifth category of cortical GABAergic cell – *MEIS2+* cells – which in mouse reside in deep layer white matter and make long-range projections. However, consistent with other reports^5^, this type was inconsistently observed across individual animals and regions, likely due to its laminar location and low abundance, and was not analyzed further.)

For each species and each of the four main classes, transcripts were pooled across cells, normalized by total number of transcripts, and scaled to 100k transcripts, which yielded four vectors of representative gene expression for each class. We then applied a series of filters to search all expressed genes for those that were selectively expressed by at least one of the four cell types in at least one species. Genes with low expression (<10 transcripts per 100k in any species) were removed. At least one species had to show a >3-fold difference between the maximum and minimum expression level across the four types. These filters identified an initial set of putative markers in one or more species. To search for genes that were consistent or differed across species, for each gene (only one-to-one orthologues were considered), Pearson correlations between pairs of species were computed, yielding six correlation values. If a gene was not detected in a given species, values were set to 0 for pairs that included that species.

### Quantitative expression level comparisons across neocortical classes

To examine the extent to which evolution has constrained a gene’s quantitative expression level across the four main neocortical cell classes, we focused on meaningfully expressed genes (4051 genes that exhibited at least 1.5-fold expression variation across the four classes in at least one species, and were also present with an abundance of at least 10 transcripts per 100k in at least one cell class). This gene list was intersected with a set of genes predicted to be intolerant of loss of function (pLI > 0.9) across 60,706 humans^6^, yielding a set of 1286 genes that were meaningfully expressed in interneurons and showed evidence of intolerance to protein-truncating variants. Pearson correlations were computed on the vector of expression values for each of these genes in all possible pairs of species.

### Species comparisons of differential expression amongst pairs of cell types

Integrated species analyses were performed using LIGER^4^ between species pairs, which had the advantage of allowing each pair of species to jointly determine cluster definition. Parameter values were explored over a range using LIGER functions to suggest optimal values; the resultant clusterings (e.g. Extended Data Fig. 2) used the following parameters: variance threshold = 0.15 (for inclusion of genes into LIGER alignment), k = 25 (number of factors), lambda = 5 (regularization parameter to penalize dataset-specific influence on alignment), resolution = 0.8 (controls resolution of clusters in community detection). For each cluster, the expression values of meaningfully expressed genes (at least 10 transcripts per 100k in both species) were extracted and fold differences for each gene was computed relative to each other cluster for each of the species independently. These fold differences were then correlated across pairs of species.

### Neocortical regionally differentially expressed genes (rDEGs)

To examine gene expression variation across neocortical regions in marmoset, region (n=7) datasets were pooled into a region-integrated LIGER analysis (variance threshold = 0.15, k = 25, lambda = 5, resolution = 0.8). These parameters produced 17 clusters; two clusters were removed for having fewer than 50 cells from one or more regions, yielding in a final set of 15 clusters for cross-region comparisons. For each cluster, differential expression was computed between all region pairs using a fold-difference threshold of 3.

#### Interneuron rDEGs in astrocytes

Marmoset neocortical astrocytes were analyzed by identifying the cluster(s) that expressed known astrocyte markers (e.g. *AQP4, GFAP, GJA1, GLUL*) from the same individuals used in interneuron analyses. Cells in these clusters were isolated from raw data and clustered using the ICA pipeline described above, which resulted in three astrocyte subtypes. For each astrocyte subcluster, fold differences of rDEGs identified in interneurons (in comparisons of PFC to V1) were computed for PFC and V1 in astrocytes.

#### Marmoset interneuron rDEGs profiled in macaque, human and mouse

Marmoset interneuron rDEGs (identified in comparisons of PFC to V1) were profiled in other species: for each rDEG, fold differences between frontal/prefrontal cortex and V1 cells were computed for each cluster identified by each species’ ICA-based clustering. The percentage and median differential expression (log_10_-transformed fold differences) of genes that were rDEGs in marmoset and were also rDEG was calculated in each species.

#### Spatial correlations

For each cluster, expression levels of rDEGs identified in comparisons between PFC and V1 were examined in the other neocortical regions (n = 5). To quantify the existence of spatial gradients, for each gene the Pearson correlation between expression and spatial order along the anterior– posterior axis was computed in the five remaining regions in order (Temp, S1, A1, Par, V2). Mean correlation values were compared to a null distribution obtained by permuting the ordering (120 possible orderings).

### Hierarchical clustering

Dendrograms of cell-type relationships were produced using hierarchical clustering (using the hclust function, method = complete, in R) of genes normalized (to 100k transcripts), using log_10_ transformed values of all expressed genes (genes with at least 10 transcripts per 100k transcripts).

### Single-molecule fluorescent *in situ* hybridization (smFISH)

#### Neocortex and hippocampus

Frozen, unfixed tissue sections (12 µm) of mouse (P60-P70; Charles River, C57BL/6; n = 2), marmoset (n = 2), and ferret (n = 1) brain tissue were cut on a cryostat (Leica CM 1950), adhered to SuperFrost Plus microscope slides (Fisher Scientific, 12-550-15) and processed for three-color smFISH using the ACD v2 RNAscope multiplexed fluorescence protocol for fresh frozen tissue. Probes are listed in Extended Data Table 2. The ferret (*Mustela putorius furo*) was sourced from Marshall Bioresources, and was used according to protocols approved by IACUC of Boston Children’s Hospital.

#### In situ–based quantification of neocortical & hippocampal LAMP5+ subtypes

For hippocampus and neocortical area S1, laminar boundaries were identified with DAPI stains in mouse and marmoset tissue. Neocortical laminae were separated into five bins (layer 1, layer 2/3, layer 4, layer 5, layer 6); hippocampal laminae within CA1 and CA2 regions were separated into four bins. Within each bin, *GAD1*+, *GAD1+/LAMP5*+, and *GAD1+/LAMP5+/LHX6*+ cells were counted in two sections of each replicate. In total, 3998 cells were counted.

#### Single-molecule FISH (smFISH) in marmoset striatum

One male marmoset (age = 6 years) was euthanized and perfused with ice-cold saline. The whole brain was immediately removed, embedded in Optimal Cutting Temperature (OCT) freezing medium, and flash-frozen in an isopropyl ethanol-dry ice bath. Samples were cut into 16 µm sections on a cryostat (Leica CM 1850), adhered to SuperFrost Plus microscope slides (Fisher Scientific, 12-550-15), and stored at −80°C until use. Samples were immediately fixed in 4% paraformaldehyde and stained on the slide according to the Advanced Cell Diagnostics RNAscope Multiplex Fluorescent Reagent Kit v2 Assay (ACD, 323100) protocol. Samples were stained for *VIP* (ACD, 554571-C2) and *NKX2-1* (ACD, 532751-C3) with antisense probes, and coverslipped with Vectashield HardSet Antifade mounting medium with DAPI (Vector Laboratories, H-1500). Z-stack serial images were taken through the whole depth on a Nikon Ti Eclipse inverted microscope with an Andor CSU-W1 confocal spinning disc unit and an Andor DU-888 EMCCD using a 20×, 0.75 NA air objective, and later max-projected in FIJI (ImageJ, NIH). Fields of view were randomly chosen across the whole striatal sample. Probes listed in Extended Data Table 2.

### Fate mapping of Lamp5+/Lhx6+ cells in mouse hippocampus and neocortex

To label *Lamp5+/Lhx6+* cells in the mouse neocortex and hippocampus, we utilized an intersectional genetics approach. In mouse mature cortical interneurons, *Id2* and *Lamp5* are expressed in nearly identical populations^7^, and so we utilized an Id2-CreER driver line ^8^ (Jax stock# 016222) in combination with an Nkx2.1-Flpo driver^9^ (Jax stock# 028577) and the Cre/Flp dependent tdTomato reporter Ai65^10^ (Jax stock# 021875) to obtain selective labeling of Lamp5/Lhx6 cells. Tamoxifen (20 mg/ml in corn oil) was administered to *Id2-CreER; Nkx2.1-Flpo; Ai65* animals (3 × 5 mg by oral gavage over 5 days) between P30–P40 to activate the CreER, after which animals were either perfused with 4% PFA/PBS and their brains processed for immunohistochemistry (20-µm cryosections; tdTomato signal was enhanced using rabbit anti-RFP from Rockland Immunochemicals; cat# 600-401-379), or acute brain slices were prepared for morphological fills as described previously^11^. Fluorescence images were acquired on a Zeiss Axio Imager.A1 and levels/contrast adjustments performed using Photoshop (Adobe).

### Ferret 10X Chromium Single Cell 3’ v3

A dataset of 801 interneurons was generated from ferret (P42) striatum. Single-nuclei suspensions from frozen tissue was generated as for Drop-seq; GEM generation and library preparation followed protocol #CG000183_ChromiumSingleCell3’_v3_UG_Rev-A. Sequencing, alignment and clustering (using the ICA pipeline described above) proceeded as for the Drop-seq datasets above.

